# Serum Alpha-2-Macroglobulin as an intrinsic radioprotective factor in patients undergoing thoracic radiation therapy

**DOI:** 10.1101/656090

**Authors:** Donata von Reibnitz, Ellen D. Yorke, Jung Hun Oh, Aditya P. Apte, Jie Yang, Hai Pham, Maria Thor, Abraham J. Wu, Martin Fleisher, Emily Gelb, Joseph O. Deasy, Andreas Rimner

**Affiliations:** Department of Radiation Oncology, Memorial Sloan Kettering Cancer Center, New York, NY; Department of Medical Physics, Memorial Sloan Kettering Cancer Center, New York, NY; Department of Laboratory Medicine, Memorial Sloan Kettering Cancer Center, New York, NY

## Abstract

**Objective:** To investigate the impact of alpha-2-macroglobulin (A2M), a suspected intrinsic radioprotectant, on radiation pneumonitis and esophagitis. Additionally, we establish multifactorial predictive models for pneumonitis and esophagitis.

**Materials/Methods:** Baseline A2M levels were obtained for 258 patients prior to thoracic radiotherapy (RT). Dose-volume characteristics were extracted from treatment plans. Spearman’s correlation (Rs) test was used to correlate A2M levels, smoking status and dosimetric variables with toxicities. Esophagitis and pneumonitis prediction models were built using least absolute shrinkage and selection operator (LASSO) logistic regression on 1000 bootstrapped datasets. Models were built using 2/3 of the data for training and 1/3 for validation.

**Results:** There were 36 (14.0%) patients with grade ≥2 pneumonitis and 61 (23.6%) with grade ≥2 esophagitis. The median A2M level was 191 mg/dL (range: 94-511). Never/former/current smoker status was 47 (18.2%)/179 (69.4%)/32 (12.4%). We found a significant correlation between baseline A2M levels and esophagitis (Rs=-0.18/*p*=0.003) and between A2M and smoking status (former or current) (Rs=0.13/*p*=0.04) but not between A2M and pneumonitis. On univariate analysis, significant parameters for grade ≥2 esophagitis included number of fractions (Rs=0.47/*p*<0.0001), treatment days (Rs=0.44/*p*<0.0001), chemotherapy use (Rs=0.40/*p*<0.0001), dose per fraction (Rs=-0.34/*p*<0.0001), total dose (Rs=0.29/*p*<0.0001), age (Rs=-0.22/*p*=0.0003), and several dosimetric variables in esophagus with Rs>0.5 (*p*<0.0001). For pneumonitis, significant clinical parameters were treatment days (Rs=0.24/*p*=0.0001), chemotherapy use (Rs=0.22/*p*=0.0004), number of fractions (Rs=0.21/*p*=0.0007), dose per fraction (Rs=-0.18/*p*=0.0035), and total dose (Rs=0.15/*p*=0.013). The most significant dosimetric variable in lung and heart was D70 (Rs=0.28/*p*<0.0001) and max dose (Rs=0.27/*p*<0.0001), respectively. LASSO bootstrap logistic regression models on the validation data resulted in the area under the receiver operating characteristic curve of 0.84 and 0.75 for esophagitis and pneumonitis, respectively.

**Conclusion:** Our findings show an association of higher A2M values with lower risk of radiation esophagitis and smoking status. Multivariate predictive models also confirmed a role of heart dose in the risk of pneumonitis.

## Introduction

Advances in radiation technology like intensity modulated radiation therapy (IMRT) and image guided radiation therapy (IGRT) have facilitated improved sparing of healthy surrounding tissues and organs when developing treatment plans. Nonetheless, radiation pneumonitis and esophagitis remain the two most common dose-limiting toxicities in thoracic RT.^1–6^ The development of either toxicity is known to depend on dosimetric and clinical variables. Concurrent chemoradiation also significantly increases the risk of developing pneumonitis or esophagitis compared to radiation alone.^7,8^

The reported incidence of radiation pneumonitis after definitive thoracic RT ranges from 10 to 20%, although figures can vary greatly.^3,9–13^ This is partly due to the fact that radiation pneumonitis remains a clinical diagnosis; there are no biomarkers or radiological findings that unequivocally confirm its presence. While many patients display only mild radiological or clinical symptoms (cough/dyspnea), which is defined as grade 1 pneumonitis per Common Terminology Criteria for Adverse Events (CTCAE) v4.03 [Supplementary Material 1], medical intervention is required with grade 2 and higher pneumonitis. Pneumonitis, which typically develops around 2 to 6 months after RT, can lead to fatal outcomes in severe cases. Severe esophageal toxicity (grade 3 to 5) occurs in around 4% of patients with sequential chemotherapy and RT and in 18-22% of patients that receive concurrent chemoradiation.^5,14^ Most patients experience mild symptoms like dysphagia or the feeling of food being “stuck” when swallowing while still undergoing radiation. Commonly opioids are needed to control dysphagia and odynophagia, but in the most severe cases, tube feeding or surgical intervention can be necessary [Supplementary Material 1].

To reduce dose-limiting toxicity in thoracic radiation, efforts have been made to adhere to normal tissue constraints derived from dose volume correlations with clinical toxicities.^15^ However, dose volume histograms do not fully predict clinical toxicities, as great interindividual variation remains. Intrinsic predictors of normal tissue radiation response may explain the variation, and improving our understanding of these is a critical component to further optimize the therapeutic ratio of thoracic RT.

Radioprotective agents, both natural and synthetic, can present an alternative method to prevent radiation-induced toxicity. Although this has been an active field of research for decades, only two compounds, amifostine and palifermine, were FDA-approved for the use in radiation therapy and neither is being commonly used in routine thoracic RT.

Another compound under investigation as an intrinsic radioprotector is alpha-2-Macroglobulin (A2M). Human A2M is a glycoprotein and the largest non-immunoglobulin serum protein. In animal studies, A2M has been shown to exhibit radioprotective effects in healthy irradiated tissue. In studies with rats that underwent full body irradiation to 6.7 Gy, rats with endo- or exogenously increased levels of A2M had a higher rate of survival, regained their baseline body weight and lymphocyte count faster, and displayed normal proliferative ability of the liver tissue compared to the control groups receiving no pre-treatment (i.e. A2M) in which all the aforementioned factors were decreased.^16–18^ Suggested key mechanisms supporting the potential of A2M as a radioprotector include its ability to promote expression of antioxidant enzymes like superoxide dismutases (SODs), inhibition of activation of fibroblasts to myofibroblasts thus preventing fibrosis, inactivation of pro-inflammatory cytokines, inhibition of all classes of proteases to maintain homeostasis, and enhancement of DNA and cell repair mechanisms by binding cytokines and growth factors.^19^ Our previous study in a small cohort showed a correlation of A2M with radiation pneumonitis.^9^ Smoking can potentially increase A2M levels however, literature specifically on A2M in smokers remains rare. Some studies confirmed higher A2M levels in smokers compared to non-smokers.^20–22^

We therefore investigated whether pre-treatment serum A2M levels are an independent predictive variable for the development of post-radiation toxicity in the lung and esophagus in a large cohort of patients receiving thoracic RT.

## Materials and Methods

### Patients

We prospectively collected data of all patients at our institution who received thoracic RT for primary malignancies or metastases at our institution between 2012 and 2016 and who had measured pre-treatment A2M values tested under an IRB waiver (n=258). Patients with any prior thoracic RT were excluded. Patients were treated with conventionally fractionated RT using 3D conformal RT (3DCRT), intensity-modulated RT (IMRT), or stereotactic body RT (SBRT). We obtained patient and treatment characteristics, smoking history, toxicity and follow-up data. Toxicity data consist of radiation pneumonitis and esophagitis rates graded per CTCAE v4.03. Data were obtained at baseline and at routine follow-up visits every three months for the first two years.

### Alpha-2-Macroglobulin

Serum samples were taken ≤30 days prior to RT start. CLIA (Clinical Laboratory Improvement Amendments) approved A2M testing was performed at Quest Diagnostics Nichols Institute (San Juan Capistrano, CA). A2M levels were given in mg/dL; the normal range was defined as 100-280 mg/dL.

### Treatment plans

For patients treated before 2014, treatment plans were retrieved from our in-house planning system.^23^ From 2014 onwards, treatments were planned in the Eclipse treatment planning system (Varian Medical Systems, Palo Alto, CA). To analyze dosimetric data, treatment plans were imported to the research platform CERR (Computational Environment for Radiological Research) and recalculated.^24^ Dosimetric variables were extracted from target structures: esophagus for esophagitis and ‘lung minus gross tumor volume (GTV)’ and heart for pneumonitis. Before that, plan doses were converted to equivalent dose in 2 Gy fractions (EQD2) with *a/b* ratio of 3 for esophagus and 10 for lung minus GTV and heart. For esophagitis, one more set of dosimetric variables were extracted by dividing the dose volume histogram (DVH) in each structure by the number of treatment days. For these fractional variables, a prefix ‘f’ was added to each dosimetric variable, for example, fmax dose. A random sample of 20 patients was used for quality assurance (QA) by verifying agreement between dose-volume metrics of the original plans and those of the recalculated DVH parameters in CERR. ‘Mean heart dose’, ‘mean esophageal dose’ and ‘lungs minus GTV’ were used for QA. The maximum differences in individual dose calculations were on the order of 10 cGy and thus considered within an acceptable range.

### Statistical methods

Univariate and multivariate analyses were performed to investigate associations between radiation-induced injuries and A2M expression, clinical, and dosimetric variables. In this study, we focused on two endpoints: esophagitis and pneumonitis. Patients were categorized into two groups for each endpoint: non-toxicity (grade 0 or 1) and clinically significant toxicity (grade 2 or greater).

A Wilcoxon rank-sum test was used to find a difference in A2M expression between the two groups. Spearman’s correlation (Rs) test was used to assess associations between endpoints, Dx values (minimum dose to the volume with the x% hottest dose in the organ of interest), computed from x=5% to x=100% in intervals of 5%, mean dose, max dose, clinical variables, and A2M.

Multivariate analysis using the least absolute shrinkage and selection operator (LASSO) logistic regression was performed using features with *p*-value < 0.05 that resulted from the univariate Spearman’s correlation test. Before the multivariate analysis, to avoid variable instability due to high collinearity, Pearson’s correlation test was conducted among all variables. A cutoff of Pearson’s correlation coefficient > 0.8 was used to determine a relatively small group of variables for further LASSO modeling.

To rigorously verify model validity, the data were split into two groups (training data with 2/3 of samples and validation data with 1/3 of samples). The model building process was carried out using only the 2/3 training data. Furthermore, to examine the stability of LASSO variable selection, the model building process was conducted using a bootstrapped dataset generated from the training data. Finally, the validation data were tested on the resulting model, quantified by the area under the receiver operating characteristic curve (AUC) as a function of sensitivity and 1-specificity. The final reported results represent the average performance on the validation data for predictive models built using 1000 bootstrapped datasets.

For statistical analyses, R language (version 3.2.4), MATLAB (version 8.6.0; MathWorks. Natick, MA) and SPSS (version 24; IBM. Armonk, NY) were used.

## Results

### Patient characteristics

In total, 258 patients were eligible for analysis. Median age was 69 years (range: 25 to 93 years) and 122 (47.3%) patients were male. Most patients were former (n=179, 69.4%) or current smokers (n=32, 12.4%). 134 patients (51.9%) underwent chemotherapy in addition to RT and the median total RT dose was 5400 cGy (range: 2700 to 7400 cGy) for conventional fractionation and 5000 cGy (range: 3000 to 10400 cGy) for SBRT. The median A2M level was 191 mg/dL (range: 94 to 511 mg/dL). More details are available in Table 1.

**Table 1.** Patient characteristics

| Factor | N | % |
| --- | --- | --- |
| Age [median, range] | 69 (25-93) years |  |
| Sex |  |  |
| *Male* | 122 | 47.3 |
| *Female* | 136 | 52.7 |
| KPS [median, range] | 90 (50-100) % |  |
| Subgroups |  |  |
| *NSCLC* | 202 | 78.3 |
| *SCLC* | 17 | 6.6 |
| *Thymoma* | 8 | 3.1 |
| *Mesothelioma* | 25 | 9.7 |
| *Lung metastases (other primary)* | 6 | 2.3 |
| Smoking History |  |  |
| *Never* | 47 | 18.2 |
| *Former* | 179 | 69.4 |
| *Current* | 32 | 12.4 |
| Pack-Years (former/current smokers) [median, range] | 37 (1-204) years |  |
| Alpha-2-Macroglobulin [median, range] | 191 (94-511) mg/dL |  |
| Chemotherapy Timing |  |  |
| *Concurrent* | 60 | 23.2 |
| *Sequential* | 74 | 28.7 |
| *No chemotherapy* | 124 | 48.1 |
| RT Total Dose [median, range] |  |  |
| *Conventional RT* | 5400 (2700-7400) cGy |  |
| *SBRT* | 5000 (3000-10400) cGy |  |

### Toxicities

Forty-nine patients (19.0%) experienced grade 1, fifty-three (20.5%) grade 2 and eight (3.1%) grade 3 radiation esophagitis. No grade 4 or 5 esophagitis was observed. Median time to development of esophagitis was 0.85 months after the start of RT (range: 0.2 to 6.47 months). Grade 1 radiation pneumonitis developed in 28 patients (10.9%), grade 2 in 26 (10.1%), grade 3 in 9 (3.5%) and grade 4 in 1 patient (0.4%). No grade 5 pneumonitis was observed. Median time to development of pneumonitis was 4.73 months after the start of RT (range: 1.3 to 8.1 months).

Of the patients who developed grade ≥2 esophagitis, 8 (13.1%) were never, 43 (70.5%) former and 10 (16.4%) current smokers. Patients with grade ≥2 pneumonitis were never smokers in 9 (25%), former smokers in 24 (66.7%) and current smokers in 3 (8.3%) cases.

### Univariate analysis

#### Alpha-2-macroglobulin

A significant correlation between baseline A2M values and esophagitis was found (Rs=-0.18/*p*=0.003). Using a Wilcoxon rank-sum test, there was a significant difference of A2M serum levels between patients with grade ≤1 versus grade ≥2 esophagitis (*p* = 0.015) when all 258 patients were analyzed as shown in Table 2. Patients with grade ≥2 esophagitis showed lower baseline serum levels of A2M than those with grade 0 or 1. For radiation pneumonitis, no statistically significant difference was found between baseline A2M levels and development of radiation pneumonitis (*p* = 0.84).

**Table 2.** Comparison of mean A2M serum levels between (unit: mg/dL) grade ≤1 and grade ≥2 esophagitis and pneumonitis. The *p*-value was calculated using Wilcoxon rank-sum test.

| Toxicity |  | Grade 0 or 1 | Grade 2+ | *p*-value |
| --- | --- | --- | --- | --- |
| Esophagitis | N | 197 | 61 | 0.015 |
|  | Mean A2M | 208.9 | 190.4 |  |
| Pneumonitis | N | 222 | 36 | 0.837 |
|  | Mean A2M | 204.1 | 207.0 |  |

A trend between smoking status and A2M levels was observed. Current smokers had higher levels (217.3 mg/dl) compared to former (207.3 mg/dl) and never smokers (185.4 mg/dl), and former smokers had higher levels compared to never smokers. The A2M level had a significant correlation with a status of former and current smoker (Rs=0.13/*p* = 0.04).

#### Clinical factors

Among standard clinical variables, the following variables showed significant correlations with both grade ≥2 esophagitis and pneumonitis, respectively: use of chemotherapy (Rs=0.40/*p*<0.0001, Rs=0.22/*p*=0.0004), number of fractions (Rs=0.47/*p*<0.0001, Rs=0.21/*p*=0.0007), dose per fraction (Rs=-0.34/*p*<0.0001, Rs=-0.18/*p*=0.0035), treatment days (Rs=0.44/*p*<0.0001, Rs=0.24/*p*=0.0001), and total dose (Rs=0.29/*p*<0.0001, Rs=0.15/*p*=0.013), and age had a significant correlation with grade ≥2 esophagitis with Rs=-0.22 (*p*=0.0003).

#### Dosimetric factors

Spearman’s correlation test between Dx in esophagus and esophagitis showed that D5 through D45 had Rs>0.50 (*p*<0.0001) as shown in Figure 1A. For the fractional dose, fD35 was the highest correlated variable with Rs=0.43 (*p*<0.0001) as shown in Figure 1B.

**Figure 1.**
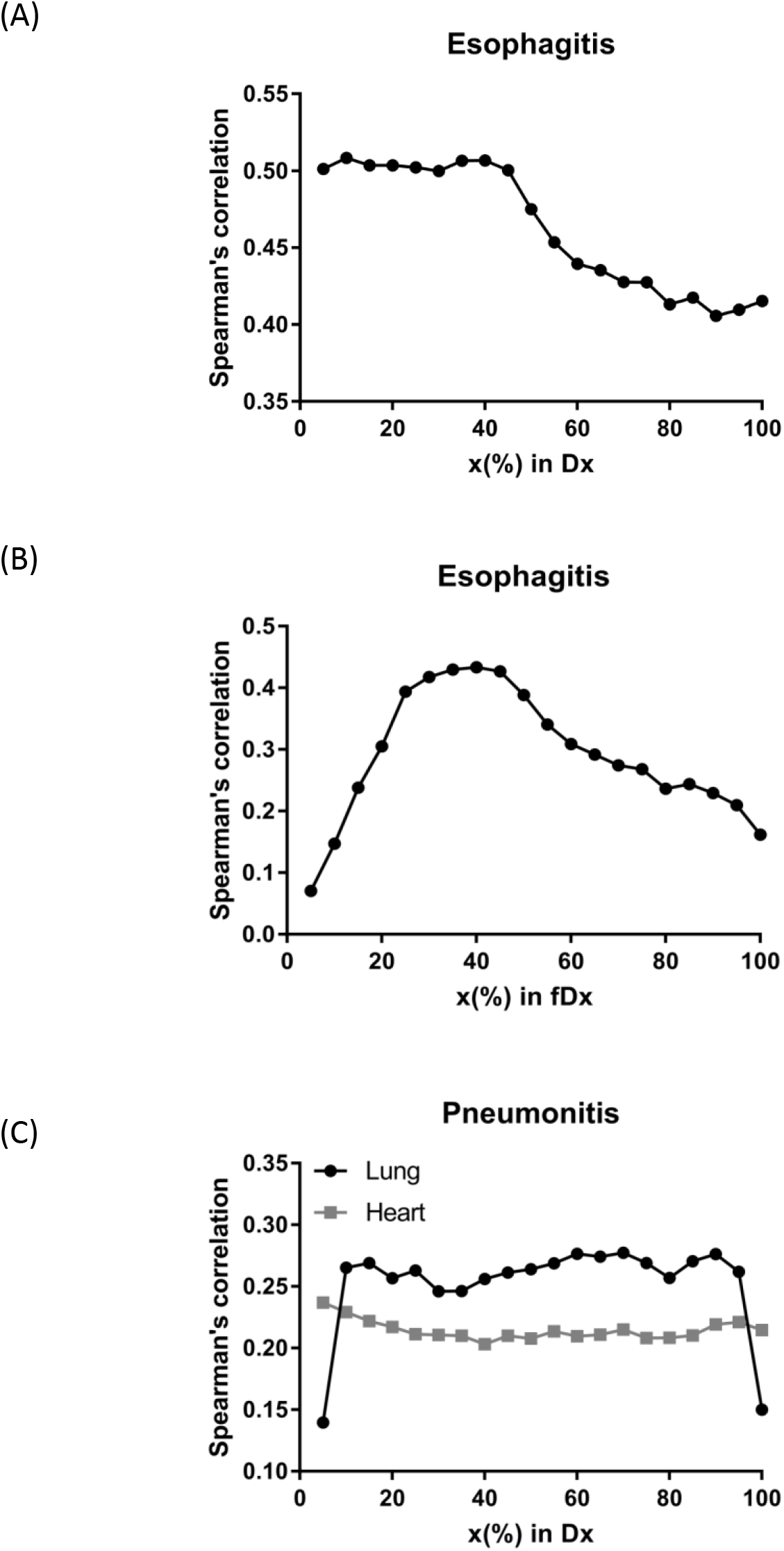
Spearman’s correlation coefficients between radiation-induced injuries (grade 2 or greater) and Dx in esophagus for (A) esophagitis, fDx in esophagus for (B) esophagitis, and Dx in lung and heart for (C) pneumonitis.

For pneumonitis, D70 (Rs=0.28/*p*<0.0001) in lung and max dose (Rs=0.27/*p*<0.0001) in heart were assessed as the highest correlated variables with pneumonitis in each organ [Figure 1C].

### Multivariate analysis and validation testing

Pearson’s correlation test using training data was performed with dosimetric variables with *p*<0.05 in the univariate Spearman’s correlation test for each organ. As can be seen in Supplementary Material 2, many dosimetric variables were highly correlated. With a threshold of 0.8 in Pearson’s correlation, redundant features were removed and one feature with the highest Rs was selected in each cluster. Clinical variables with *p*<0.05 in the univariate analysis and dosimetric variables left after Pearson’s correlation test were used in the LASSO logistic regression: (D10, D35, D40, D65, D85, fD20, fD25, fD35) in esophagus, age, and A2M for esophagitis; (D10, D15, D70, D90) in lung, (D5, D55, D95, max dose) in heart for pneumonitis; for both endpoints, treatment days, chemotherapy use, dose per fraction, total dose, and number of fractions were used. In addition, indication of SBRT treatment (coded as 1 and 0 for SBRT and non-SBRT treatment) was used as a variable.

LASSO logistic regression models were trained using bootstrapped datasets generated from training data and were tested on the validation data, resulting in an average AUC of 0.84 (standard deviation [SD]=0.03) and 0.75 (SD=0.06) for esophagitis and pneumonitis, respectively [Supplementary Material 3]. Additional modeling was performed for esophagitis without A2M which resulted in the same average AUC (0.84). This appears to be due to more significant dosimetric and clinical variables used in the modeling. To assess the importance of features, the frequency of occurrence of each feature during the model building process was counted [Figure 2]. For the esophagitis model, D85 and chemotherapy were most frequently selected with 755 and 717 times, respectively. It is worthy to note that A2M was selected with 621 times, implying its high correlation with esophagitis. Taken together, although A2M did not further improve performance of the predictive model for esophagitis, its association with esophagitis was confirmed. For the pneumonitis model, max dose in heart was most frequently selected with 792 times. Patients were sorted based on predicted outcomes on the validation data and grouped into six equal bins with 1 being the lowest risk group and 6 being the highest risk group. When comparing observed and predicted incidence, we found a high conformity of both endpoints, meaning that the predictive models are highly robust [Figure 3]. Final predictive models built using all training data are shown in Supplementary Material 4.

**Figure 2.**
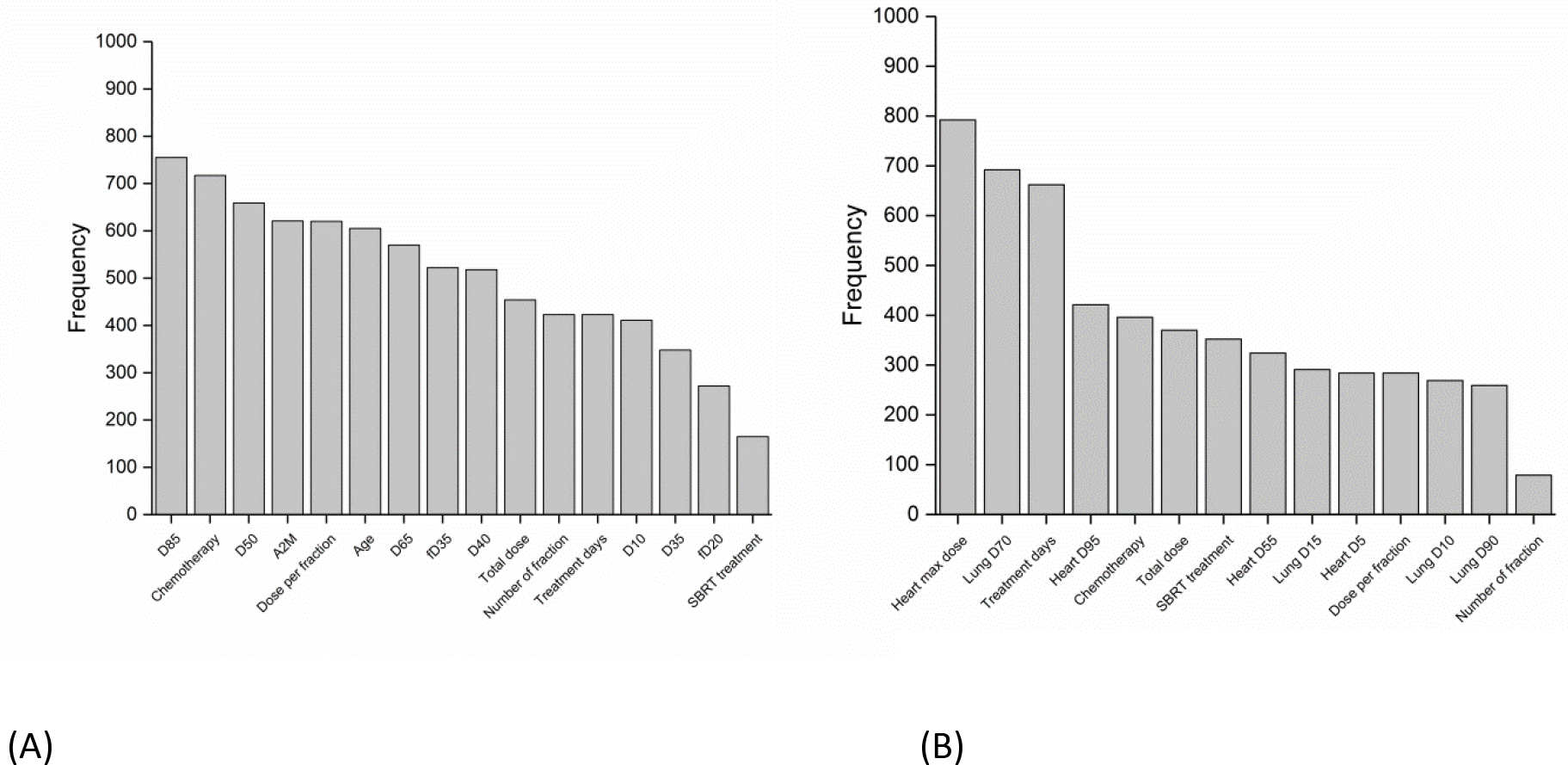
Frequency of occurrence of each feature used in 1000 predictive models for (A) esophagitis and (B) pneumonitis.

**Figure 3.**
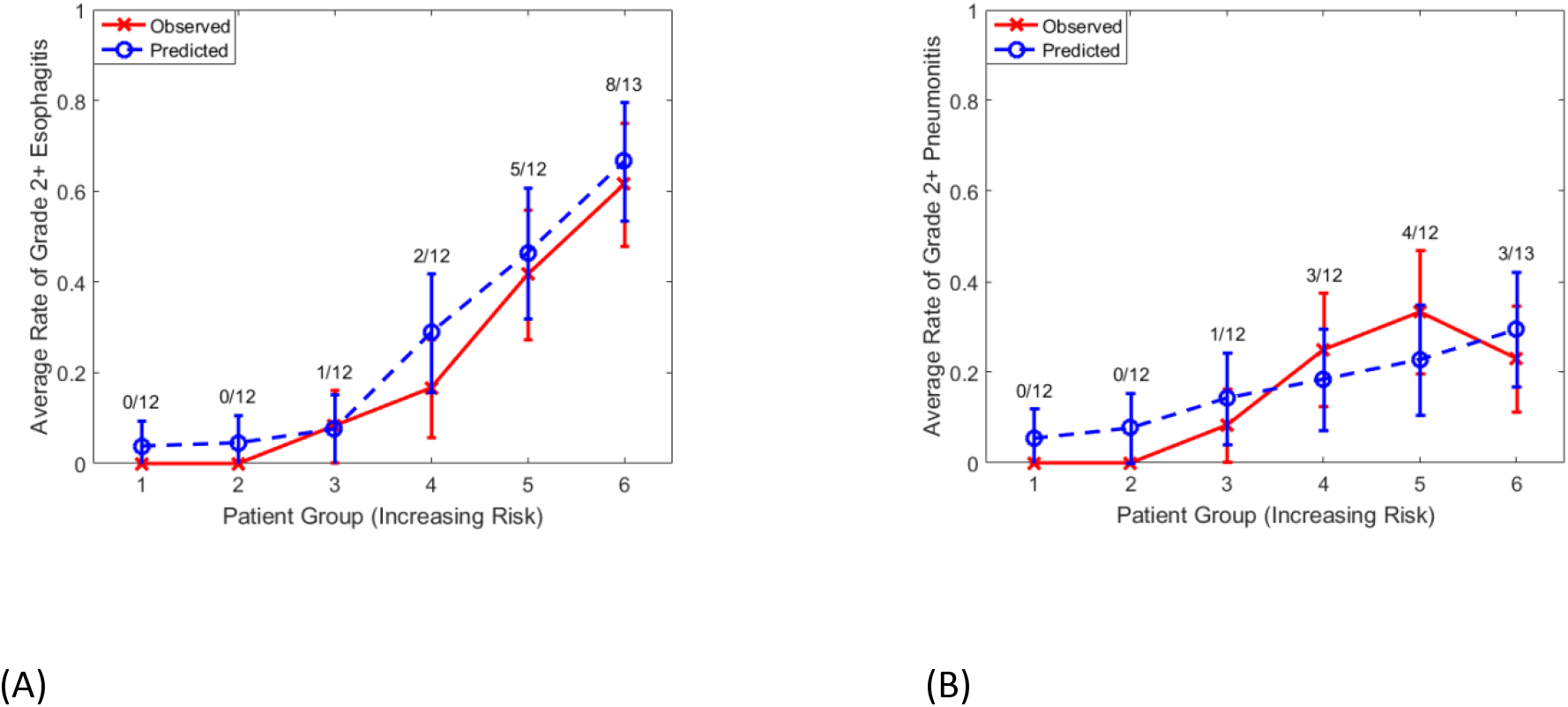
Comparison of observed and predicted incidence on validation data (1/3 of samples) for (A) esophagitis and (B) pneumonitis. The numerator and denominator indicate the number of events and the number of samples in each bin, respectively.

In addition, the frequency of occurrence of a pair of features used in the LASSO logistic regression model was investigated [Figure 4], which provides the information of interaction effects of features in the predictive model.

**Figure 4.**
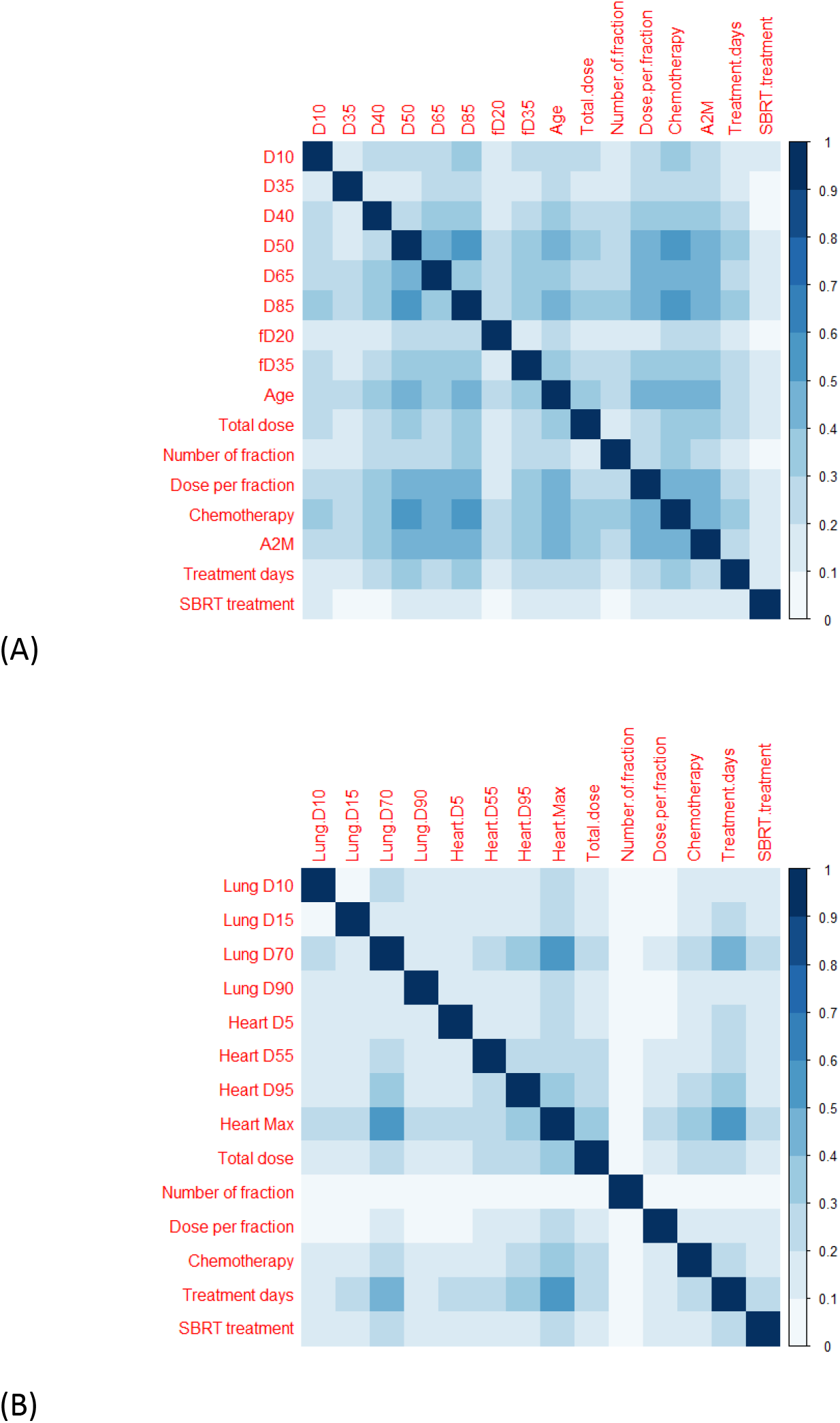
Frequency of occurrence of a pair of features used in 1000 predictive models for (A) esophagitis and (B) pneumonitis. The frequency of occurrence of a pair of features was divided by 1000.

## Discussion

Our study showed that there is an association between low natural pre-treatment baseline A2M serum levels in patients with thoracic malignancies and an increased risk of developing radiation esophagitis. This finding suggests that higher levels of A2M may have a protective effect in patients undergoing thoracic RT. This result is in line with previous reports from studies in mice.^16–18^ Although we found this association for esophagitis, we were unable to detect such a correlation between A2M and pneumonitis rate. This may be due to an effect of the uneven distribution of smoking history in the high- and low-risk toxicity groups. For example, of the patients that developed esophagitis, 13.1% (n=8) were never smokers, while there were 25% (n=9) never smokers among those who developed pneumonitis.

As we identified in our correlative analysis, A2M levels appear to be influenced by patients’ smoking status. Former and current smokers displayed higher A2M values than patients that had never smoked. The effect of smoking on the immune system has been studied extensively. Paradoxically, smoking results in immunosuppression in regard to infections as well as aggravated autoimmunity. Altered levels of inflammatory cytokines like TNF-α, IFN-γ, IL-1β, IL-6, IL-8, IL-10 and others have been reported in healthy smokers.^25–28^ A possible explanation for the connection between active smoking and a lower risk for esophagitis or pneumonitis is that long-term cigarette smoking leads to an increased immune response in the lung and surrounding tissues due to the damage it inflicts on the lung parenchyma. Although the mechanisms resulting in normal tissue injury after RT are still under investigation, the release of reactive oxygen species (ROS) as well as proinflammatory and profibrotic cytokines is thought to have a central role in the process.^29^ Higher baseline levels of acute-phase proteins like A2M may have a protective effect on the irradiated tissue by binding proinflammatory and profibrotic cytokines, thus reducing the acute cytokine toxicity, and inducing an upregulation of antioxidant enzymes like manganese superoxide dismutase (MnSOD).^19,29^

Taking into account the multifactorial etiology of radiation toxicity^29^, it is essential to look at different predictive factors in the development of lung and esophageal injury after RT. Most commonly, different dosimetric parameters are included in predictive models for pneumonitis and esophagitis but biological and genetic determinants are also under investigation.^9,11,29–36^ In our analysis, we focused on dose-volume metrics, age, chemotherapy, and other clinical variables in addition to A2M. The high selection frequency of A2M in the model building process also confirms our primary correlative analyses linking A2M levels to esophagitis rates. Factors that have repeatedly shown significant correlation with esophagitis include V40-V60^6,37–40^ (Vx: percentage volume receiving at least x Gy) and the mean esophageal dose ^41–43^. Several authors have reported on the increased risk for high-grade esophagitis after sequential and especially concurrent chemo-RT in comparison to RT alone.^5,6,44–47^

For pneumonitis, we were able to validate the correlation with radiation dose received by the heart (in our model, max dose in heart). Different lung dose volumes (V5-V40 and mean dose in lung) have been found to predict the development of pneumonitis^7,10,48^. In addition, the dose received by the heart during thoracic radiation seems to be an accurate predictor.^49,50^ The best fitting predictive model reported by Huang et al. included the following variables, in order of selection: D10 in heart, D35 in lung and max dose in lung, and had an AUC of 0.72.^49^ Although the ideal dosimetric variable(s) for predicting pneumonitis across all patient subgroups may not yet be known, it is evident that heart doses are an essential part of any model built for this cause. Though we could not confirm the impact of A2M on pneumonitis with our data, a correlation between them has been previously described.^9,50^ We may have been limited by the lower incidence of grade ≥2 pneumonitis (14.0%) and the low rate of current smokers in our patient cohort. In the previously published study on A2M and pneumonitis, pneumonitis rates were between 19 and 35%. Furthermore, we confirmed chemotherapy in conjunction with RT (either concurrent or sequential) as a significant factor for the development of pneumonitis, evident by its fifth-highest frequency in 1000 model runs [Figure 4B], consistent with multiple other studies.^7,48^

While all patients had A2M prospectively collected before their RT and toxicity data were systematically prospectively scored per CTCAE v4.03, caution is warranted regarding the interpretation of these results. In particular, including a factor like chemotherapy in predictive models should be considered carefully as different regimens, doses and timings, depending on the patient population, make it a very heterogeneous variable. Similarly, the patient cohort we studied was diverse in regard to diagnosis and treatment. Although requirements for eligibility included no prior RT, patients underwent different modes of RT (conventional, SBRT, IMRT) which may have an impact on the toxicity profile. Furthermore, as noted in the beginning of the discussion, smoking status as a variable was not evenly distributed. Given its proposed link to serum A2M levels, this could be a cause for certain discrepancies in our results.

## Conclusion

In summary, the analysis of our institutional dataset has produced predictive models for both esophagitis and pneumonitis. This is the first report on the apparent protective function of higher A2M levels in regard to esophagitis. While using A2M is untested as a prophylactic radioprotective agent in humans at the present time, our study provides an incentive to utilize pre-treatment A2M levels as an indicator for the potential risk of developing radiation-induced toxicity. However, further independent validation and clinical trials are needed to establish A2M as a dependable biomarker and potential radioprotective mediator.

## Acknowledgements

This research was funded in part through the NIH/NCI Cancer Center Support Grant P30 CA008748.

## Supplementary Material

**Supplementary Material 1.**
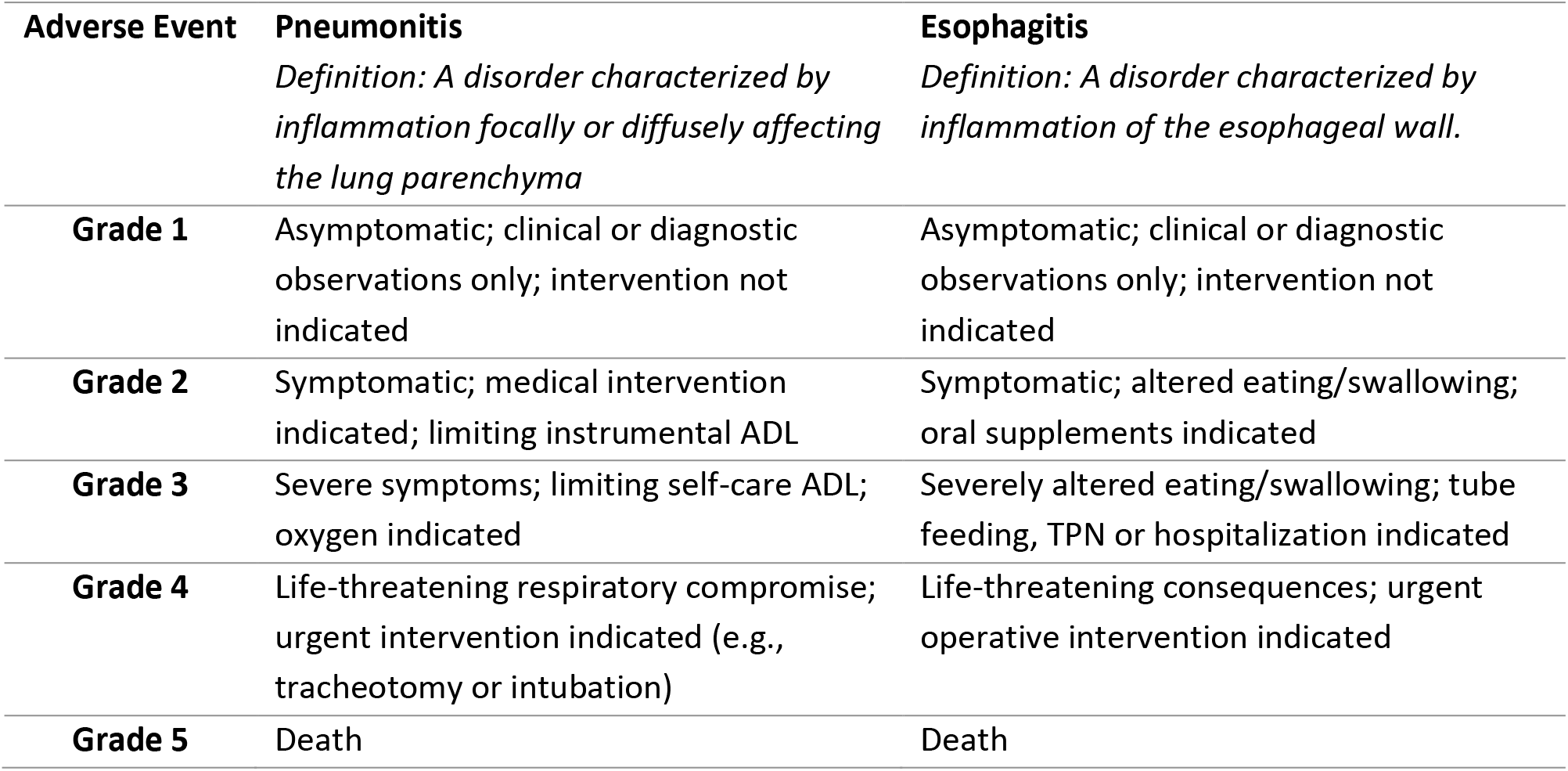
CTCAE v4.03 grading for pneumonitis and esophagitis.

**Supplementary Material 2.**
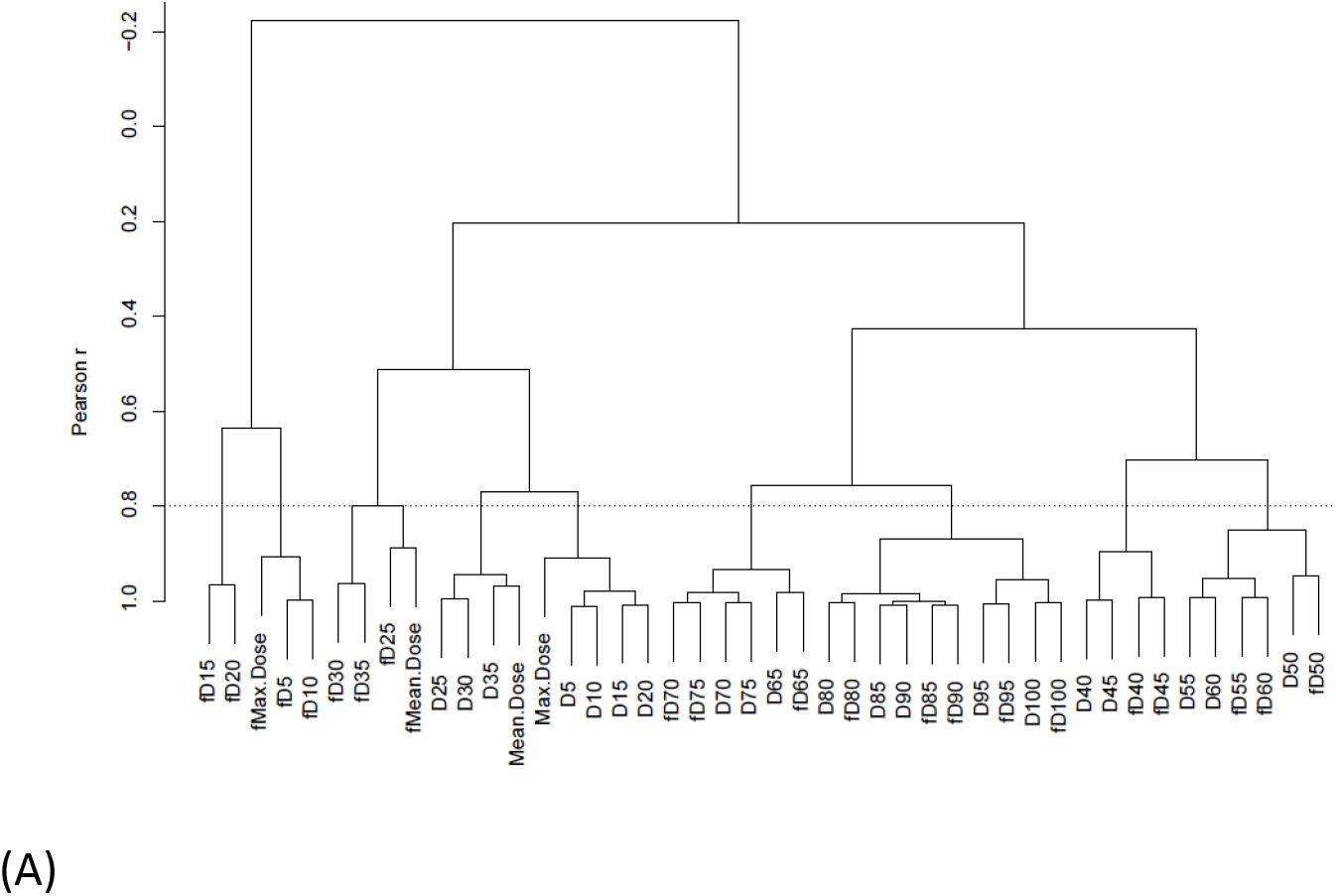
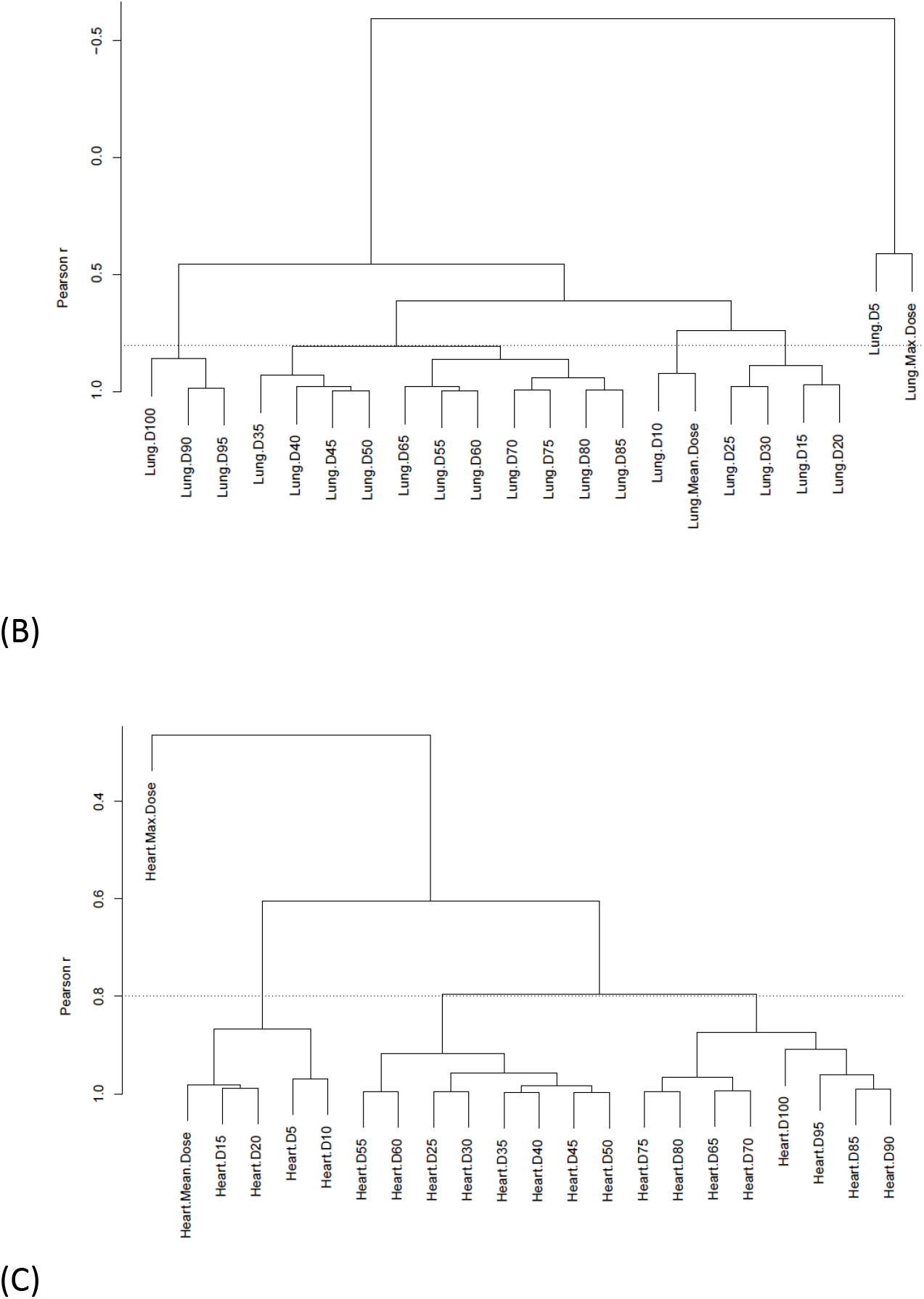
Pearson’s correlation test using training data to remove redundant features with a threshold of 0.8: (A) dosimetric variables in esophagus for esophagitis, (B) in lung for pneumonitis, and (C) in heart for pneumonitis.

**Supplementary Material 3.**
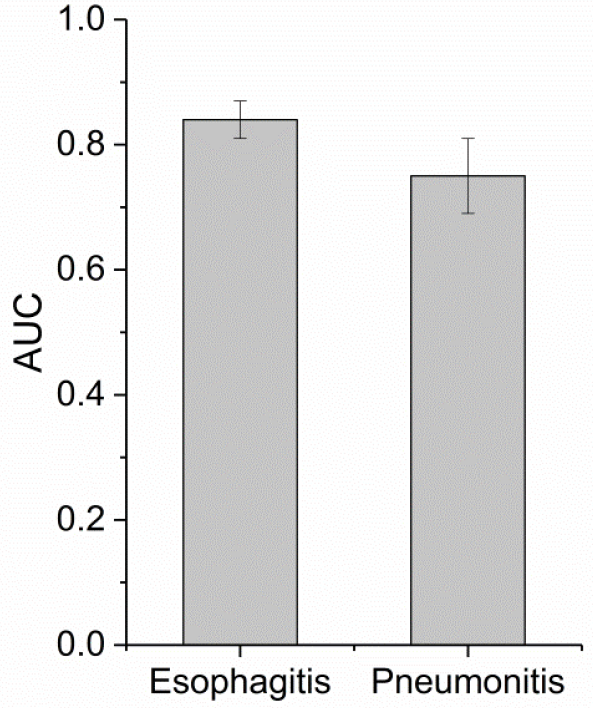
Average AUC after 1000 iterations of the LASSO logistic regression modeling on the validation data. The error bar indicates the standard deviation.

**Supplementary Material 4.**
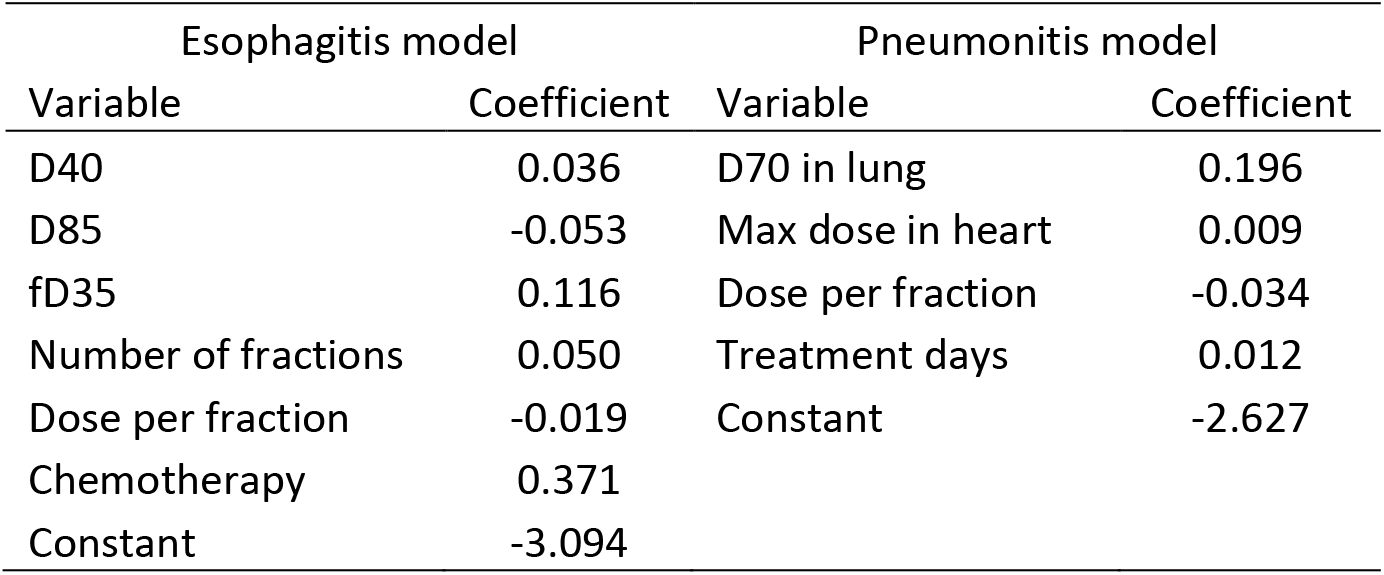
Final predictive models for esophagitis and pneumonitis.

